# Multimodal optical mesoscopy reveals the quantity and spatial distribution of gram-positive biofilms in *ex vivo* tonsils

**DOI:** 10.1101/2023.07.03.547470

**Authors:** Megan Clapperton, Tash Kunanandam, Catalina D. Florea, Catriona M. Douglas, Gail McConnell

**Affiliations:** Department of Physics, SUPA, University of Strathclyde, Glasgow, UK; Department of Otolaryngology – Head and Neck Surgery, Royal Hospital for Children, Glasgow, UK; Department of Otolaryngology – Head and Neck Surgery, Queen Elizabeth University Hospital, Glasgow, UK; Strathclyde Institute of Pharmacy and Biomedical Sciences, University of Strathclyde, Glasgow, UK

**Keywords:** tonsil, biofilms, mesoscopy, multimodal, reflection, fluorescence, image analysis

## Abstract

Biofilms are known to be present in tonsils, but little is known about their spatial location and size distribution throughout the tonsil. Studies of the location and distribution of biofilms in tonsil specimens have thus far been limited to either high-magnification methods such as electron microscopy, which enables high resolution imaging but only from a tiny tissue volume, or lower magnification techniques such as light microscopy, which allow imaging of larger specimens but with poor spatial resolution. To overcome these limitations, we report the use of multimodal optical mesoscopy to visualize and quantify the number and spatial distribution of gram-positive biofilms in fresh, excised paediatric tonsils. This methodology supports simultaneous imaging of both the tonsil host and biofilms in whole mounts of tissue up to 5 mm × 5 mm × 3 mm with subcellular resolution throughout. A quantitative assessment of thirty-six tonsil specimens revealed no statistically significant difference between biofilm presence on the tonsil surface and the interior of the tonsil. This new quantitative mesoscale imaging approach may prove useful in understanding the role of biofilms in tonsillar diseases and other infections.

## 1. Introduction

Antibiotics are routinely prescribed to patients presenting with tonsillar diseases such as acute recurrent tonsillitis with the aim of eradicating bacterial infection [1]. However, tonsillar diseases are becoming more resistant to antibiotics due to the persistence of bacteria through the formation of biofilms. Biofilms are communities of bacteria encased in a protective matrix, and they are known to be more resistant to antibiotic treatment than planktonic bacteria [2].

Imaging studies have confirmed the presence of biofilms in tonsil tissue [3], but data are limited because tonsils are too large to study with a conventional microscope. In children (0 – 16 years), mean tonsil volume is 1.5 ± 0.9 cm^3^ [4]. Imaging data are therefore either representative of only a small volume of the relatively large tonsil specimen or they show results from much larger tissue volumes but without the spatial resolution needed for imaging of biofilms.

At small spatial scales, Chole and Faddis [3] used a combination of light microscopy and transmission electron microscopy in combination with absorption contrast agents to study biofilms in fixed tonsil specimens. Their work showed strong anatomical evidence for the presence of biofilms in chronically diseased tonsils, but their method was limited, visualising only a single biofilm with light microscopy at high optical magnification. Also, the absorption contrast staining methods used in their work rely on chemical fixation of tissues prior to imaging, which causes tissue shrinkage and distortion of both host and bacterial structures that can complicate morphometric analysis [5], [6]. Diaz et al [7] evaluated the performance of both absorption staining and fluorescence staining using brightfield transmission microscopy and confocal laser scanning microscopy for the study of biofilms in pediatric tonsillar disease. Their findings suggest that biofilms are present in pathological tonsils, as with the work of Chole and Faddis [3], imaging was performed at high optical magnification (100x) with additional digital magnification (resulting in images of up to 400x) and imaging was only performed within very tiny sub-regions of the whole tonsil. Quantitative studies of biofilms in tonsils have been performed with light microscopy and electron microscopy methods but the small imaging volumes possible with these methods have thus far offered only limited insights into the size or location of biofilms in the tissue [3], [8], [9].

At larger spatial scales, optical coherence tomography (OCT) has been used to image excised human tonsils and has proven successful for identifying different tonsil tissue components including epithelium, dense connective tissue, lymphoid nodules, and crypts [10]. This method relied on scattering and tissue autofluorescence for low-contrast imaging of structures, and features such as biofilms could not be discriminated from the tonsil host. Additionally, although imaging of tissues several millimetres in size was possible the spatial resolution of the images was considerably poorer than that possible in the light microscope studies previously discussed, with a reported resolution of tens of microns. Another technique offering a large imaging volume is magnetic resonance imaging (MRI) [11], [12]. However, the resolution of conventional clinical MRI is around 1-2 mm [13], which is too poor to resolve small biofilms. Specialized high-resolution MRI setups, such as that of Herrmann et al [14], have increased the resolution but only to around 100 μm. MRI solutions offer poorer resolution than OCT, but both techniques lack the molecular specificity required to identify biofilms in tissue.

In sum, little is currently known about the spatial extent and location of biofilms in tonsil tissues because of limitations in imaging technology, yet this information could prove vital in helping to identify biofilm burden and disease pathogenesis.

Optical mesoscopy is a rapidly developing field that enables visualisation of larger samples than is possible with standard light microscopy without compromising on spatial resolution [15]. In practice, optical mesoscopy involves imaging of objects ranging from millimetres up to centimetres in size with μm or nm resolution. As such, optical mesoscopy spans the boundary between classic biological imaging and preclinical ‘biomedical’ imaging, typically utilising low magnification objective lenses with a large field of view. The Mesolens is one such objective, which offers the unusual combination of low magnification (4x) with a high numerical aperture (0.47), and it can image specimens up to 5 mm × 5 mm × 3 mm in size with 0.7 μm lateral and 7 μm axial resolution throughout this entire volume [16].

Here, we describe a new multimodal confocal mesoscopy methodology for the visualization and quantitative analysis of gram-positive biofilms in whole mounts of fresh pediatric tonsil tissue up to 5 mm × 5 mm × 3 mm in size. We developed a point-scanning confocal reflection imaging mode for use with the Mesolens to visualise tonsillar tissue structure, and we have used this together with point-scanning confocal fluorescence contrast to reveal the location and scale of gram-positive biofilms in the tonsil host. We show that this methodology can reveal biofilms with sub-cellular resolution in tonsil tissue in a single three-dimensional dataset. We have also developed a simple image analysis pipeline that is compatible with the large datasets generated by the Mesolens, and we have applied this for the quantitative study of biofilms at different locations in patient tissue.

## 2. Methods

### 2.1. Specimen Preparation

This study was approved by Biorepository Management Committee of NHS Greater Glasgow and Clyde, UK (Biorep 548). Thirty-six pediatric tonsils were collected after routine tonsillectomy at the Royal Hospital for Children, Glasgow, UK. Whole tonsils were rapidly transported in sterile saline solution (0.9% Sodium Chloride, Baxter Healthcare Ltd, UK) to the Strathclyde Institute of Pharmacy and Biomedical Sciences at the University of Strathclyde, Glasgow, UK, within 2 hours of excision from the patient. Care was taken to minimise mechanical disruption to the sample by keeping it as still as possible during transport, and by using saline at an appropriate volume to only just fully submerge the tonsils. Tonsils were dissected using a fresh, sterile No. 27 surgical blade (12474070, Fisher Scientific, UK) to produce whole mounts of either the outer tonsillar surface or internal tonsillar tissue with volumes of approximately 5 mm × 5 mm × 3 mm.

Gram-positive bacteria located within tonsil tissue were labelled using a Vancomycin-BODIPY FL conjugate (V34850, ThermoFisher Scientific, USA). The stain was diluted in dH_2_O to yield a final concentration of 0.5 μg/ml. The dissected specimens were incubated in 5 ml of this working solution in 15 ml universal tubes for 60 minutes at room temperature. Specimens were covered with aluminium foil to reduce bleaching by ambient light. After staining, the specimen was gently washed without agitation in Hanks Balanced Salt Solution (HBSS, Gibco, Fisher Scientific, UK), pH 7.7, (3x, 1 min). Imaging of the tonsil specimens began within 3 hours of surgery.

### 2.2. Specimen mounting for multimodal optical mesoscopy

Individual whole mount specimens were secured in a custom-designed mount for long-term imaging with the Mesolens. A diagram of the disassembled specimen holder is shown in Figure 1. The specimen mounts were manufactured using Invar to minimise thermal expansion to the specimen holder and hence reduce any specimen movement during image acquisition. The top and bottom sections correspond to the immersion chamber. The bottom part of the chamber is comprised of an Invar plate that incorporates a glass window to form a well. The whole mount of tonsil tissue was placed into this well and was surrounded with HBSS as the imaging mountant. A 70 mm × 70 mm type 1.5 coverslip (0107999098, Marienfeld, Germany) was placed atop the specimen and bottom plate, avoiding bubbles in the HBSS mounting medium. The top plate of the specimen mount contained a 48 mm diameter hole in the top plate. The underside of this top plate incorporated a nitrile ‘O’ ring 38 mm in diameter to be held against the coverslip using screws at the exterior of the chamber. When in contact with the coverslip this arrangement created a well. After assembly, dH_2_O was added into this well, creating a stable water bath within the ring with the coverslip as its base and the front of the Mesolens dipping into it. Surface tension proved sufficient to preserve the water column (up to 3 mm high) for extended periods (>18 hours).

**Figure 1.**
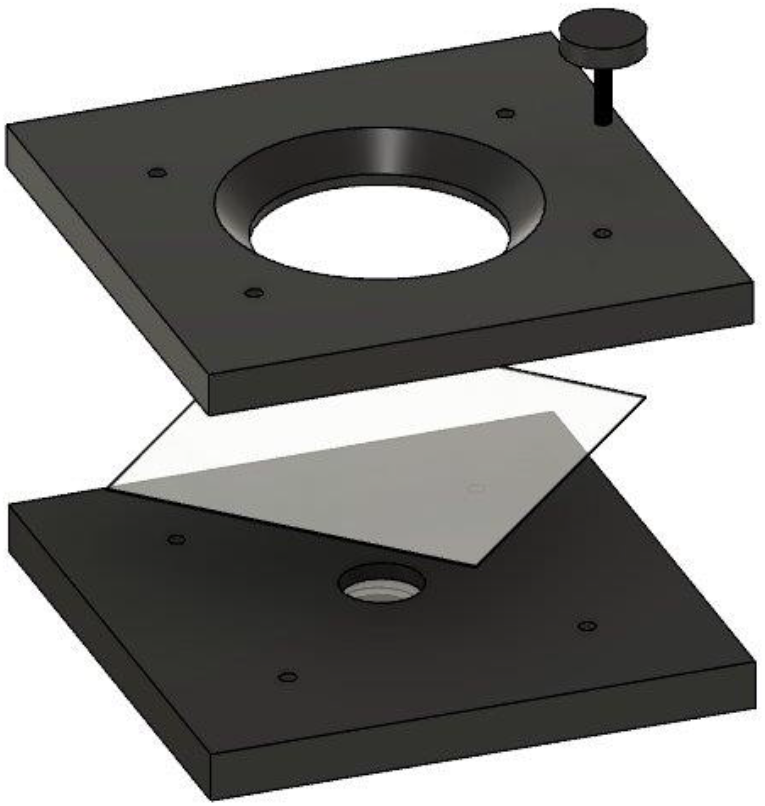
Computer-aided design drawing of the disassembled specimen holder and multipart chamber for long-term imaging of whole mounts of tonsil tissue using water immersion. The top and bottom sections correspond to the immersion chamber. The top plate includes a 38 mm diameter ‘O’ ring (not shown) that is brought into contact with the large coverslip. Screws (only one shown, though four are used in practice) bring the three sections together. Distilled water is added to the bath created by the contact of the top plate and coverslip of the specimen slide for long-term imaging.

### 2.3. Multimodal optical mesoscopy

A technical report of the Mesolens instrument has been published previously [16], therefore we report here only a brief overview of the Mesolens together with modifications to the system made specifically for multimodal imaging of biofilms in whole mounts of tonsil tissue.

A schematic of the multimodal setup using the Mesolens is shown in Figure 2. A 488 nm laser (Multiline Laserbank, Cairn Research) was used as an illumination source to produce reflection contrast images resulting from the refractive index boundary at the HBSS to tonsil interface, and the same laser was used to excite fluorescence from gram-positive bacteria labelled with Vancomycin-BODIPY FL. Reflection contrast was chosen over autofluorescence to map the tonsil tissue. Tonsil tissue is autofluorescent and has previously been used for the study of tonsil architecture [10] but the autofluorescence emission signal spectrally overlaps with the fluorescence emission of Vancomycin-BODIPY FL and therefore was not suitable for multimodal imaging of specimens in this work. The reflection signal at 488 nm was easily spectrally discriminated from the fluorescence from Vancomycin-BODIPY FL using optical filters, and this arrangement resulted in high contrast images in both reflection and fluorescence channels. A total laser power of 4 mW at the specimen plane was used for imaging of all specimens.

**Figure 2.**
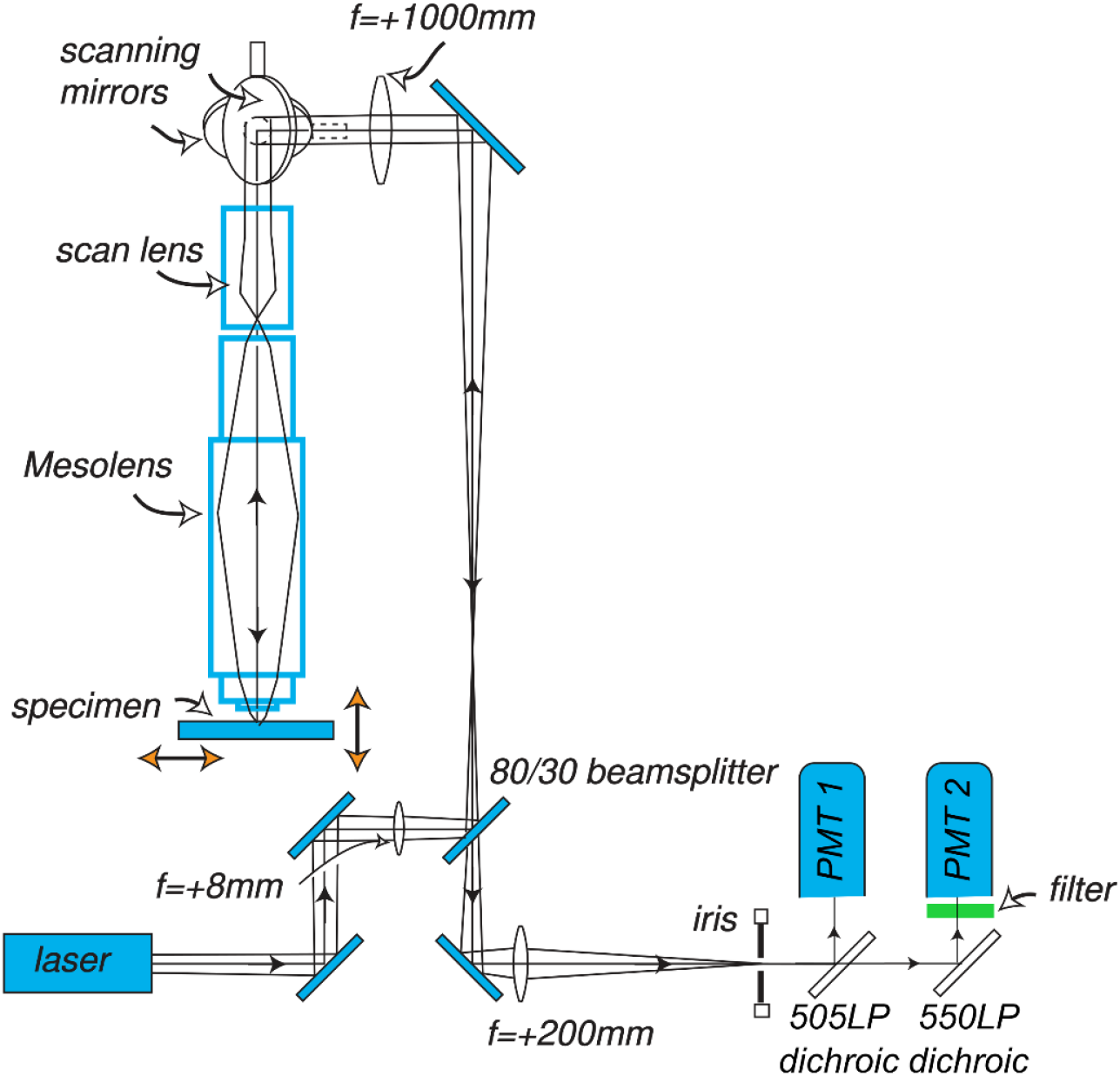
Schematic diagram of the multimodal point-scanning confocal Mesolens. A laser of 488 nm wavelength was used for simultaneous imaging with reflection contrast and fluorescence contrast. A single confocal iris was used in the detection path. Two photomultipliers (PMTs) were used for detection of the reflection (PMT 1) and fluorescence (PMT 2) contrast with 505 nm and 550 nm long pass filters, respectively. The filter in the detection path between the 505LP dichroic and PMT 2 was a 520 ± 35 nm band pass filter, and this served to block scattered laser light and transmit only fluorescence from gram-positive bacteria labelled with Vancomycin-BODIPY FL to the photodetector.

The Mesolens was used in point-scanning confocal laser scanning mode for imaging of all specimens. A 5 mm × 5 mm lateral field of view was scanned with a sampling rate of 2 pixels per micron to satisfy the Shannon-Nyquist criterion [17]. The pixel dwell time was set to 1 μs, and no frame averaging was performed. This led to an acquisition time of 100 s for a single optical section of the full 5 mm × 5 mm field.

The reflection and fluorescence signals were spectrally separated for detection using a 505 nm dichroic mirror (DMLP505R, Thorlabs, USA) that reflected only this wavelength, and transmitted longer wavelength fluorescence signals. The fluorescence signal between 505 nm and 550 nm was reflected by a second dichroic mirror (DMLP550R, Thorlabs, USA), and an additional band pass filter specified for transmission of 520 ± 35 nm (520/35 BrightLine HC, Semrock, USA) was used to reject any backscattered laser light and transmit only fluorescence. Two photomultiplier tubes (PMTs) (P30-01, Senstech, UK) were used for detection of reflection (PMT 1) and fluorescence (PMT 2).

Movement of the specimen along the optical axis for imaging at discrete z planes using the confocal method was performed using a computer-controlled z-positioning system (Optiscan II, Prior Scientific). To minimise data acquisition time the z-step size was set to between 3 μm and 7 μm, with smaller step sizes used for thinner specimens and larger step sizes used for thicker tissues. The system was controlled using an in-house, laser scanning software package, ‘Mesoscan’, designed to handle scanned images. Images were stored in the Open Microscopy Environment OME.TIFF format.

### 2.3. Image processing and analysis

A single-colour optical section of the 5 mm × 5 mm field of view of the Mesolens with lateral sampling of 2 pixels/μm corresponds to a file size of just over 190 Mb. With multimodal imaging and millimetre thick specimens, the datasets generated from these specimens ranged in size from 30 Gb to 130 Gb depending on the complexity of the tonsil specimen surface and the overall depth of imaging. Datasets were processed and analysed using a 64-bit Windows server with two intel Xeon Silver 4114 CPU processors at 2.20 and 2.19 GHz and 1.0 TB installed RAM.

Images for presentation purposes were first contrast adjusted using the Contrast Limited Adaptive Histogram Equalization (CLAHE) 15 function in FIJI [18] with the default parameters (block size = 127, histogram bins = 256, maximum slope = 3.00). FIJI was also used to create dual-channel maximum intensity projections of multimodal mesoscopy datasets that gave a two-dimensional overview of the size and location of biofilms relative to the whole mount of tonsil tissue.

To visualise and quantify data in three-dimensions, OME.TIFF files from the two channels were sequentially opened with Imaris and these data were subsequently converted to the proprietary Imaris file format (Imaris 9.8, Oxford Instruments). The ‘Surfaces’ tool was used to segment objects in each channel from the background to facilitate measurements [19]. Next, the ‘Statistics’ function was used to extract volumetric information from the image data. The volume of the tonsil imaged using reflection contrast was measured with this simple method, and no additional processing was required.

To evaluate the presence of planktonic bacteria in the tonsil specimens, we assumed a single spherical coccus of between 1 μm and 2 μm in diameter [20], and therefore we calculated the volume of a single bacterial cell to be between 0.52 μm^3^ and 4.19 μm^3^. The number of discrete object volumes within this volumetric range in the fluorescence channel were counted using the ‘Filter’ tool in Imaris and these data correspond to the number of single bacterial cells in the tonsil specimens.

To assess the total biofilm volume, based on previous work which reported biofilms to be spherical bacterial aggregates with a minimum diameter of 20 μm [21] we calculated biofilms had a volume equal to or greater than 4180 μm^3^. The number of discrete object volumes with this minimum volume in the fluorescence channel were counted using the ‘Filter’ tool in Imaris and these data corresponded to the number of biofilms in the tonsil specimens. These numerical data were extracted from Imaris and were exported to Microsoft Excel (Microsoft 365 Apps for Enterprise, Microsoft Corporation, USA) for further analysis.

To quantify the biofilm volume relative to the volume of the tonsil host, the ratio of the total biofilm volumes to the tonsil volume was calculated for each specimen from the tonsil surface (n=17) and tonsil interior (n=19) respectively. Plots of these data were generated using GraphPad Prism (GraphPad Prism v.8.0.2, GraphPad Software).

## 3. Results

Using multimodal optical mesoscopy it proved possible to visualize the presence of individual bacteria and biofilms within an unusually large volume of fresh tonsil tissue.

Figure 3(A) shows a maximum intensity axial projection of a whole mount of tonsil tissue prepared for imaging as described in Sections 2.1 and 2.2. The tonsil was imaged as described in Section 2.3, using a 5 μm z-step size over a total axial range of 810 μm. These data took just over 4.5 hours to acquire. The tonsil tissue is shown in green and gram-positive bacteria are shown in magenta. The tonsil specimen extended beyond the imaged field, but even within this 5 mm × 5 mm lateral area it is evident that biofilm spread was highly heterogeneous in this specimen. A large biofilm extending over several square millimetres is shown close to the centre of the field. Several smaller discrete biofilms are also visible, and these are distributed unevenly within the tissue. A region of interest (ROI) is highlighted with a yellow box. A digital zoom of this ROI is shown in Figure 3(B). This ROI shows the level of spatial detail that is obtained at all positions in the maximum intensity projection dataset. A biofilm is visible adjacent to hundreds of individual bacterial cells. A white dotted line in 3(B) shows a further ROI that includes both the biofilm in the centre of the image and adjacent individual bacteria. The intensity of the magenta channel only is plotted in Figure 3(C). These data confirm that the diameter of this biofilm is approximately 25 μm and that individual bacteria between 1 μm to 2 μm in size are clearly resolved. These data demonstrate the suitability of multimodal optical mesoscopy to simultaneously visualize individual bacteria and biofilms over extended spatial volumes in a single dataset.

**Figure 3.**
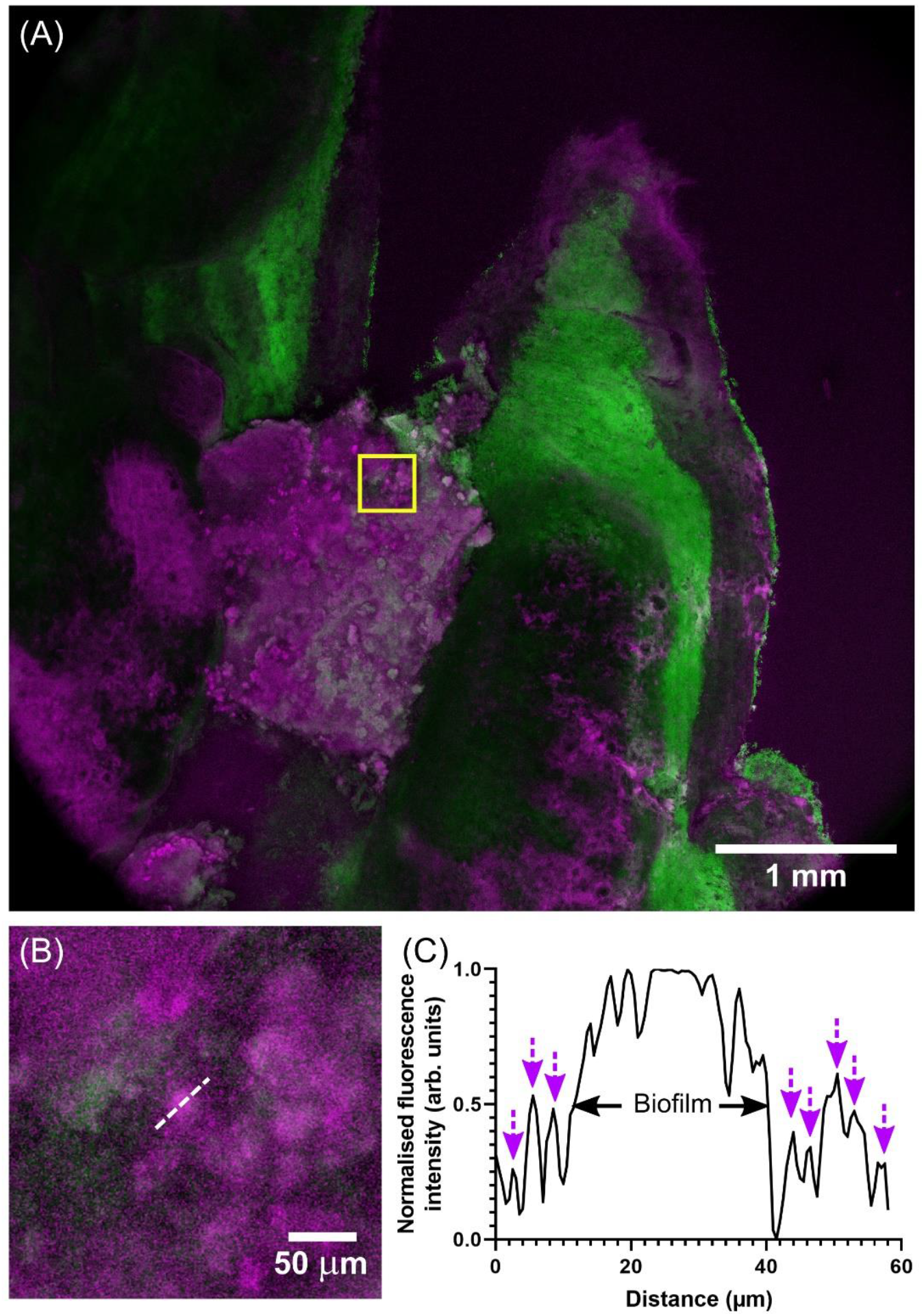
Multimodal optical mesoscopy provides an extended view of large biofilms in whole mounts of fresh tonsil with sub-cellular resolution. (A) Maximum intensity projection of a whole mount of fresh tonsil. The tonsil specimen was imaged over a 5 mm × 5 mm 810 μm volume using a z-step size of 5 μm. The tonsil tissue is shown in green, and biofilms are shown in magenta. Biofilms are unevenly distributed throughout the tissue volume. A yellow box shows a tiny region of interest. (B) Digital zoom of the region highlighted with the yellow box in 3(A). Several biofilms are surrounded by individual bacteria. A white dotted line highlights a sub-ROI through the centre of one biofilm. (C) Line intensity profile of the magenta channel only from the white dotted line ROI indicated in 3(B). The biofilm is measured to be around 25 μm in diameter and it is surrounded by individual bacteria of between 1 μm to 2 μm in size. The position of bacteria along this line ROI are indicated by magenta arrows.

Figure 4 shows a three-dimensional render of a different tonsil specimen to that presented in Figure 3, imaged as per the methods reported in Sections 2.1 and 2.2. These data were obtained over a 5 mm × 5 mm × 1 mm imaging volume with a z-step size of 5 μm, which took just over 5.5 hours to acquire. A video showing this dataset from different angles and at various levels of display zoom is available as Movie 1. In these data the complex three-dimensional topography of the tonsil surface (shown in green) is clearly visible with voids in the tissue corresponding to the opening of crypts. An example of a visible crypt is highlighted with a cyan arrow. The highly heterogeneous spatial distribution of gram-positive bacteria (shown in magenta) is clearly visible in three-dimensions. Without further digital zoom individual bacteria are not visible, but they are present and can be seen in the raw data (available for download). However, individual biofilms are present at this level of display zoom, both across the tissue surface and at the opening of tonsil crypts. An example of biofilms surrounding the mouth of a crypt is shown with a yellow arrow. Throughout the tissue biofilms appear as bright punctate magenta objects on the order of 20 μm to 200 μm in diameter. We note that these thick biofilms are frequently connected to thinner biofilms, which are more widely spread across the tissue surface. There are only a few regions where the green and magenta signals spatially coincide: this co-occurrence is visible as white areas. From this low coincidence we concluded that biofilms are mostly adhering to the tonsil surface in this specimen, rather than being embedded within the tissue.

**Figure 4.**
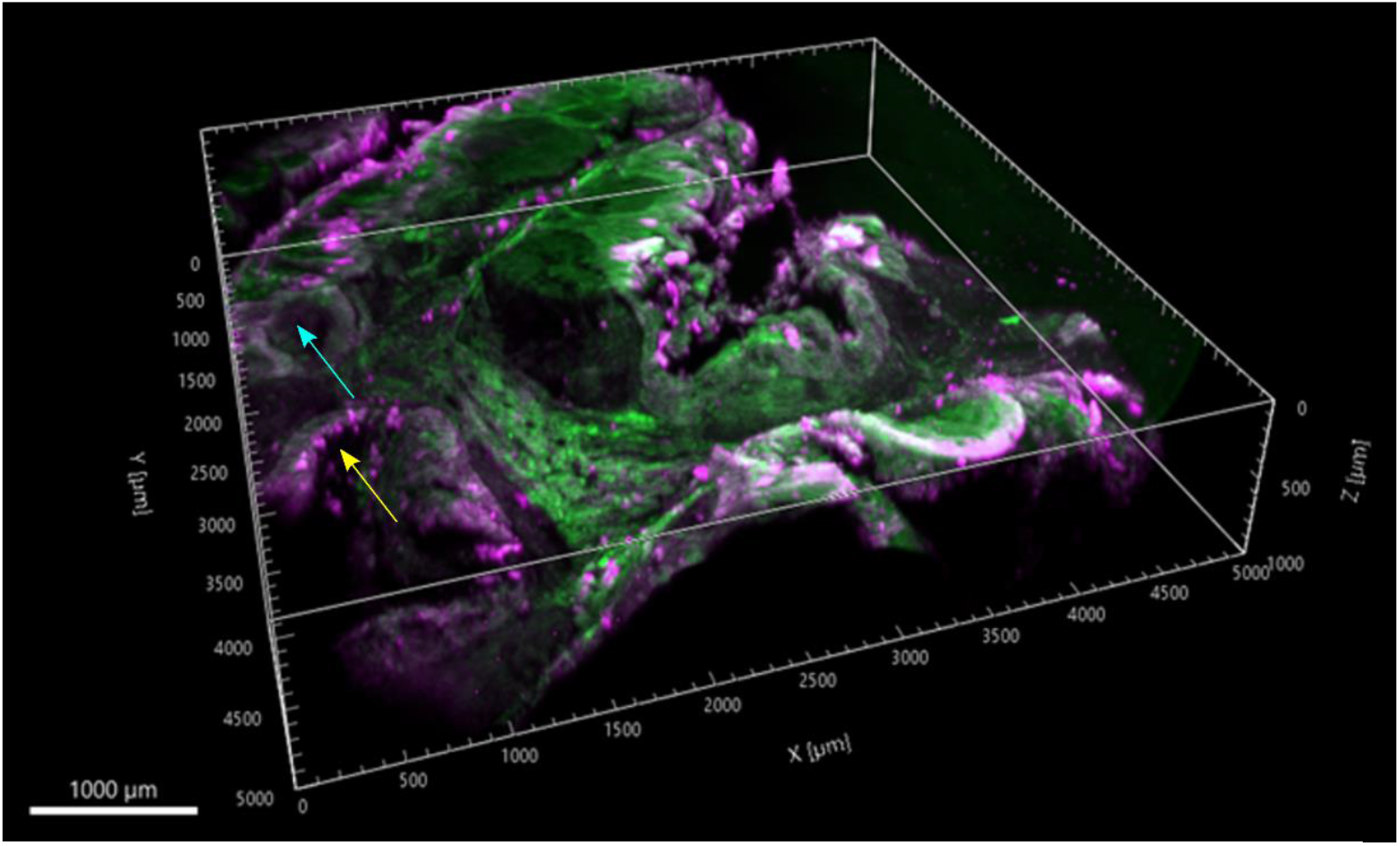
Three-dimensional render of a whole mount of fresh *ex vivo* tonsil. Tonsil tissue is shown in green, and biofilms are shown in magenta. The image volume is 5 mm × 5 mm × 1 mm, and this was obtained with a z-step size of 5 μm. The complex tonsil tissue architecture is clearly visible, with voids in the tissue corresponding to the opening of crypts, an example of which is highlighted with a cyan arrow. The spatial distribution of biofilms is heterogeneous, with biofilms of various shapes and sizes extending over the tonsil surface and lining the entrance to crypts: an example region showing biofilms surrounding the mouth of a crypt is shown with a yellow arrow.

We analysed the data from n=36 specimens (n=17 tonsil surface, n=19 tonsil interior) as described using the methods in Section 2.3. Results are shown in Figure 5. As Figure 5(A) shows, biofilms were detected in all image datasets, with the minimum and maximum number of biofilms observed per sample being 58 and 1229, respectively. A larger number of gram-positive biofilms was measured in whole mounts of tissue from the tonsil surface with a mean of 481 biofilms compared to tissue from the tonsil interior with a mean of 339 biofilms. Data were not statistically significant. Figure 5(B) shows the ratios of total biofilm volume to tonsil volume for image datasets obtained from the tonsil surface and tonsil interior. Specimens with both a high (close to 1.5:1) and low (0.001:1) biofilm volume were measured, with similar mean biofilm: tonsil ratios obtained for both specimens from the tonsil surface and tonsil interior. These values were measured as 0.32:1 and 0.35:1, respectively. We interpret these data to mean that irrespective of location in the tonsil the ratio of gram-positive biofilms to tonsil tissue is broadly consistent.

**Figure 5.**
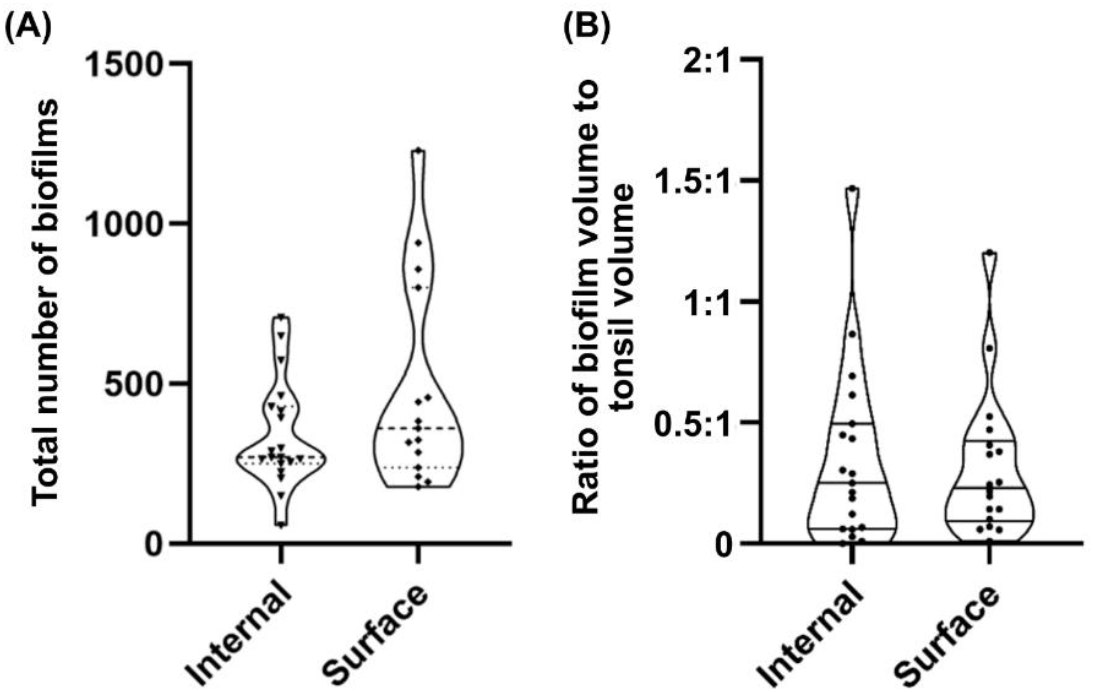
Quantitative analysis of biofilms in whole mounts of fresh tonsil. (A) Measured total number of biofilms in whole mounts of fresh tonsil. Biofilms were observed in all specimens. A larger number of gram-positive biofilms was measured in whole mounts of tissue from the tonsil surface with a mean of 481 biofilms compared to tissue from the tonsil interior with a mean of 339 biofilms. Data were not statistically significant. (B) shows the ratios of total biofilm volume to tonsil volume for image datasets obtained from the tonsil surface and tonsil interior. Similar biofilm: tonsil volume ratios were obtained for both specimens from both regions, with mean values of as 0.32:1 and 0.35:1 for the surface and interior, respectively.

## Discussion

We have demonstrated multimodal optical mesoscopy for the visualization and analysis of biofilms *in situ* in multi-millimetre-sized whole mounts of fresh, *ex vivo* tonsil tissue with subcellular resolution. Our results show that biofilms were present in all n=36 specimens imaged. This result is to the contrary of other studies including the work of Chole and Faddis [3] which have reported smaller positive outcomes. We attribute our findings to the greatly increased imaging volume with high spatial resolution that is possible with the Mesolens compared to other imaging methods. If we assume imaging of a whole mount of tonsil tissue using a conventional point-scanning confocal microscope equipped with a 20x objective this gives a calculated imaging volume of approximately 700 μm × 700 μm × 500 μm. By comparison, the Mesolens can image a specimen that is more than 300-times greater in volume. This increase in imaging volume considerably increases the probability of detecting biofilms – or even single bacteria – in these large specimens.

Our analysis reveals little difference in the number of biofilms present in on the surface and interior of tonsil tissues, but the number of biofilms varies considerably between different specimens, with the maximum number of biofilms observed being 20x that of the specimen with the fewest biofilms present. Again, this measurement can only be made at present using multimodal optical mesoscopy.

In this work we have focused exclusively on the study of gram-positive biofilms, but our methodology could be adapted to include the visualization and analysis of gram-negative biofilms in the tonsil host. For example, fluorescent Bocillin 650/665 could be used to label both gram-negative and gram-positive bacteria, and the Vancomycin-BODIPY FL conjugate we have used could be used as a specific label for gram-positive bacteria only. The difference in peak fluorescence emission wavelength between these two labels is over 100 nm, and hence they would be compatible for simultaneous labelling and multimodal imaging. Another possibility is the use of fluorescence in situ hybridisation (FISH) for the direct visualisation of specific bacterial species. FISH has previously been applied for simultaneous imaging of five bacterial species in microscopic volumes of tissue [22]. However, while multimodal optical mesoscopy with FISH would enable species-level study of much larger specimens, FISH requires fixation and, as described previously, this may cause tissue distortion. It would also be possible to use an additional fluorescent label for the study of the tonsil architecture rather than using reflection contrast but given the large volume of the whole mount specimens this would soon become costly for imaging of more than a small number of specimens.

Although there are powerful open-source software tools such as BiofilmQ [23] and BASIN [24] for the analysis of biofilm image data, the large file sizes of the Mesolens data are not immediately compatible with these resources. We also explored the utility of image analysis tools with general applications in biology and biomedicine such as DeProj [25], but we encountered similar difficulties in working with our large file sizes. We used Imaris in this work for rendering and analysis of our image datasets, but we recognize the potential of napari [26] as an open-source alternative for future multimodal optical mesoscopy studies.

The standard treatment for acute recurrent tonsillitis is antibiotic medication but reports suggest that biofilms in tonsils may be eradicated by wiping the tonsil surface to mechanically disrupt biofilms [27]. While the complex architecture of tonsils includes crypts that serve as reservoirs for biofilms [3], our work confirms that biofilms can also be large and widely spread across the tonsil surface, covering areas extending many square millimetres in size. This new observation may prove useful in developing new cleaning protocols and antimicrobials that target biofilms on the tonsil surface.

We have already used the Mesolens in point-scanning confocal fluorescence mode to study whole mature colony biofilms [28, 29], but this work reports the first application of the Mesolens to the study of fresh, *ex vivo*, human tissue. With proof-of-principle now established, this newly developed multimodal optical mesoscopy imaging method can be readily applied to biofilm research related to other areas of human health including diseases of the middle ear such as otitis media with effusion and wounds in skin [30].

## Acknowledgements

This work was supported by the Glasgow Children’s Hospital Charity, grant number GCHCRF/PHD/2020/02, UK Research and Innovation, grant numbers BB/P02565X/1, BB/T011602/1, BB/W019032/1 and MR/K015583/1, and the Scottish Universities Life Sciences Alliance. C.M.D. was partly supported by the Chief Scientific Office and UK Research and Innovation, grant number MR/W030381/1. G.M. was partly supported by the Leverhulme Trust.

## Author Contributions

G.M. and C.D. conceived the experiments, defined the methodology, acquired funding, and supervised and administered the project. M.C. performed the investigation, including data curation, visualization, and formal analysis. C.F. and T.K. provided resources. M.C. and G.M. wrote the original draft, and M.C, G.M, C.D and C.F were involved in reviewing and editing.

## Data Availability statement

The data generated and analysed during the study are available upon reasonable request from the corresponding author.

## Competing interests

The authors declare no competing interests.

## Supplementary information

https://strath-my.sharepoint.com/:v:/g/personal/g_mcconnell_strath_ac_uk/ER0U4hc6Zk9IrWL7Y2jR9g0BWc7gEWYBlCn_7_h4d2B8Tw?e=ExWO9k (no login required)

Movie 1. Three-dimensional render of a whole mount of fresh tonsil of the data shown in Figure 4. Tonsil tissue is shown in green, and biofilms are shown in magenta. The image volume is 5 mm × 5 mm × 1 mm and this was obtained with a z-step size of 5 μm. The complex tonsil tissue architecture is clearly visible, with voids in the tissue corresponding to the opening of crypts. The spatial distribution of biofilms is heterogeneous, with biofilms of various shapes and sizes extending over the tonsil surface and lining the entrance to crypts.

## Notes

https://strath-my.sharepoint.com/:v:/g/personal/g_mcconnell_strath_ac_uk/ER0U4hc6Zk9IrWL7Y2jR9g0BWc7gEWYBlCn_7_h4d2B8Tw?e=ExWO9k

### Competing Interest Statement

The authors have declared no competing interest.

## References

[1] Munck, H, Jørgensen, W. W., & Klug, T. E. Antibiotics for recurrent acute pharyngo-tonsillitis: systematic review. Eur. J. Clin. Microb. Inf. Dis. 37, 1221–1230 (2018).

[2] Patel, H. H., Straight, C. E., Lehman, E. B., Tanner, M., & Carr, M. M. Indications for tonsillectomy: A 10 year retrospective review. Int. J. Pediatr. Otorhinolaryngol. 78, 2151–2155 (2014).

[3] Chole, R. A. & Faddis, B. T. Anatomical Evidence of Microbial Biofilms in Tonsillar Tissues: A Possible Mechanism to Explain Chronicity. Arch. Otolaryngol. Neck Surg. 129, 634–636 (2003).

[4] Michaels, L. Normal Anatomy, Histology; Inflammatory Diseases. Ear, Nose and Throat Histopath. Springer, 265–272 (1997) ISBN 9781447133322.

[5] Swidinski, A. et al. Spatial organisation of microbiota in quiescent adenoiditis and tonsillitis. J. Clin. Pathol. 60, 253–260 (2006).

[6] Mazzone, R. W., Kornblau, S. & Durand, C. M. Shrinkage of lung after chemical fixation for analysis of pulmonary structure-function relations. J. Appl. Physiol. 48, 382–385 (1980).

[7] Diaz, R. R., et al., Relevance of biofilms in pediatric tonsillar disease. Eur. J. Clin. Microbiol. Infect. Dis. 30, 1503–1509 (2011).

[8] Woo, J. H., Kim, S. T., Kang, I. G., Lee, J. H., Cha, H. E. & Kim, D. Y. Comparison of tonsillar biofilms between patients with recurrent tonsillitis and a control group. Acta Otolaryngol. 132, 1115–1120 (2012).

[9] Alasil, S. M., Omar, R., Ismail, S., Yusof, M. Y., Dhabaan, G. N., & Abdulla, M. A. Evidence of Bacterial Biofilms among Infected and Hypertrophied Tonsils in Correlation with the Microbiology, Histopathology, and Clinical Symptoms of Tonsillar Diseases. Int. J. Otolaryngol. 2013, 1–11 (2013).

[10] Pahlevaninezhad, H., et al., Optical Coherence Tomography and Autofluorescence Imaging of Human Tonsil. PLoS ONE 9, e115889 (2014).

[11] Suto, Y. Matsuda, E. & Inoue, Y. MRI of the pharynx in young patients with sleep disordered breathing. Br. J. Radiol. 69, 1000–1004 (1996).

[12] King, A. D., Lei, K. I. K., & Ahuja, A. T. MRI of primary non-Hodgkin’s lymphoma of the palatine tonsil. Br. J. Radiol. 74, 879, 226–229 (2001).

[13] Lin, E., & Alessio, A. What are the basic concepts of temporal, contrast, and spatial resolution in cardiac CT? J. Cardiovasc. Comput. Tomogr. 3, 403–408 (2009).

[14] Herrmann, K.-H., et al., High-resolution MRI of the human palatine tonsil and its schematic anatomic 3D reconstruction. J. Anat. 240, 166–171 (2022).

[15] Munck, S., et al. Challenges and advances in optical 3D mesoscale imaging. J. Microsc. 286, 201–219 (2022).

[16] McConnell, G., Trägårdh, J., Amor, R., Dempster, J., Reid, E., & Amos, W. B. A novel optical microscope for imaging large embryos and tissue volumes with sub-cellular resolution throughout. eLife 5, e18659 (2016).

[17] Shannon, C. E. Communication in the Presence of Noise. Proc. IRE 37, 10–21 (1949).

[18] Schindelin, J., et al. Fiji: an open-source platform for biological-image analysis. Nat. Meth. 9, 676–682 (2012).

[19] Battistella, E., Schniete, J., Wesencraft, K., Quintana, J. F., & McConnell, G. Light-sheet mesoscopy with the Mesolens provides fast sub-cellular resolution imaging throughout large tissue volumes. iScience 25, 104797 (2022).

[20] Balaji, K., Thenmozhi, R., & Pandian, S. K. Effect of subinhibitory concentrations of fluoroquinolones on biofilm production by clinical isolates of Streptococcus pyogenes. Ind. J. Med. Res. 137, 963–9711 (2013).

[21] Roberts, A. L., et al. Detection of group A Streptococcus in tonsils from pediatric patients reveals high rate of asymptomatic streptococcal carriage. BMC Pediatr. 12, 3 (2012).

[22] Clark, S. T., Waldvogel-Thurlow, S., Wagner MacKenzie, B., Douglas, R. G., & Biswas, K. Fluorescence in situ hybridisation in Carnoy’s fixed tonsil tissue. Sci. Rep. 12, 12395 (2022).

[23] Hartmann, R., et al. Quantitative image analysis of microbial communities with BiofilmQ. Nat. Micro. 6, 151–156 (2021).

[24] Hartman, T., et al. BASIN: A semi-automatic workflow, with machine learning segmentation, for objective statistical analysis of biomedical and biofilm image datasets. J. Mol. Biol. 435, 167895 (2023).

[25] Herbert, S., et al. LocalZProjector and DeProj: a toolbox for local 2D projection and accurate morphometrics of large 3D microscopy images. BMC Biol. 19, 136 (2021).

[26] Sofronieuw, N., et al. napari: a multi-dimensional image viewer for Python. 10.5281/zenodo.3555620.

[27] Ciftci, Z., Develioglu, O., Arbak, S., Ozdoganoglu, T., & Gultekin, E. A new horizon in the treatment of biofilm-associated tonsillitis. Ther. Adv. Respir. Dis. 8, 78–83 (2014).

[28] Rooney, L. M., Amos, W. B., Hoskisson, P. A., & McConnell, G. Intra-colony channels in E. coli function as a nutrient uptake system. ISME J. 14, 2461–2473 (2020).

[29] Bottura, B., Rooney, L. M., Hoskisson, P. A., & McConnell, G. Intra-colony channel morphology in Escherichia coli biofilms is governed by nutrient availability and substrate stiffness. Biofilm 26, 100084 (2022).

[30] Vestby, L. K., Grønseth, T., Simm, R., & Nesse, L. L. Bacterial biofilm and its role in the pathogenesis of disease. Antibiotics 9, 59 (2020).

